# Extreme gradient boosting machine learning algorithm identifies genome-wide genetic variants in prostate cancer risk prediction

**DOI:** 10.1101/2023.10.27.564373

**Authors:** David Enoma, Victor Chukwudi Osamor, Ogunlana Olubanke

## Abstract

Genome-wide association studies (GWAS) identify the variants (Single Nucleotide polymorphisms) associated with a disease phenotype within populations. These genetic differences are essential in variations in incidence and mortalities, especially for Prostate cancer in the African population. Given the complexity of cancer, it is imperative to identify the variants that contribute to the development of the disease. The standard univariate analysis employed in GWAS may not capture the non-linear additive interactions between variants, which might affect the risk of developing Prostate cancer. This is because the interactions in complex diseases such as prostate cancer are usually non-linear and would benefit from a non-linear Machine Learning gradient boosting viz XGBoost (extreme gradient boosting). We applied the XGBoost algorithm and an iterative SNP selection algorithm to find the top features (SNPs) that best predict the risk of developing prostate cancer with a Support Vector Machine (SVM). The number of subjects was 907, and input features were 1,798,727 after appropriate quality control. The algorithm involved ten trials of 5-fold cross-validation to optimize the dataset’s hyperparameters and the prediction task’s second module (utilizing SVM). The model achieved AUC-ROC cure of 0.66, 0.57 and 0.55 on the Train, Dev and Test sets, respectively. The area under the Precision-Recall Curve was 0.69, 0.60 and 0.57 on the Train, Dev and Test sets, respectively. Furthermore, the final number of predictive risk variants was 2798, associated with 847 Ensembl genes. Interaction analysis showed that Nodes were 339 and the edges were 622 in the gene interaction network. This shows evidence that the non-linear Machine learning approach offers excellent possibilities for understanding the genetic basis of complex diseases.

## 1. Introduction

Genome-wide association studies are conducted to elucidate genetic variants (SNPs-Single Nucleotide Polymorphisms) within a population that confer disease risk [1]. This approach has helped identify the significance of risk variants that affect the etiology of the disease. Moreover, it does not consider the non-linear interaction of SNPs that may have smaller effect sizes because of the stringent p-values enforced by multiple testing Fields[2].

More suitable approaches, such as non-linear Machine Learning methods, will help capture non-linear interaction among SNPs to identify SNPs of greater significance to the disease in question [3]. Furthermore, the problem in GWAS datasets is a problem with many features and a smaller sample size [4]. This means there are many SNPs and a significantly smaller number of samples. This makes it relevant to evaluate the importance of variables applied to a model.

This work aimed to apply a modified extreme gradient boosting (XGBoost) algorithm [5] to Prostate cancer GWAS data. XGBoost was used to select SNPs in a breast cancer risk prediction task. It is based on gradient-boosted decision trees, which, unlike penalized regression approaches, may contain complicated non-linear interactions in a non-additive form in prediction models [6]. XGBoost can capture non-linear interactions between the SNPs. The linear SVM was then used to distinguish cancer cases and healthy controls.

The modified XGBoost algorithm by [5] is an iterative SNP selection algorithm that finds the most predictive SNPs of developing prostate cancer in the target population. Figure 1 below details the algorithmic approach, which captures the non-linear interactions to find the best SNPs that are the predictive prostate cancer phenotype.

**Figure 1.**
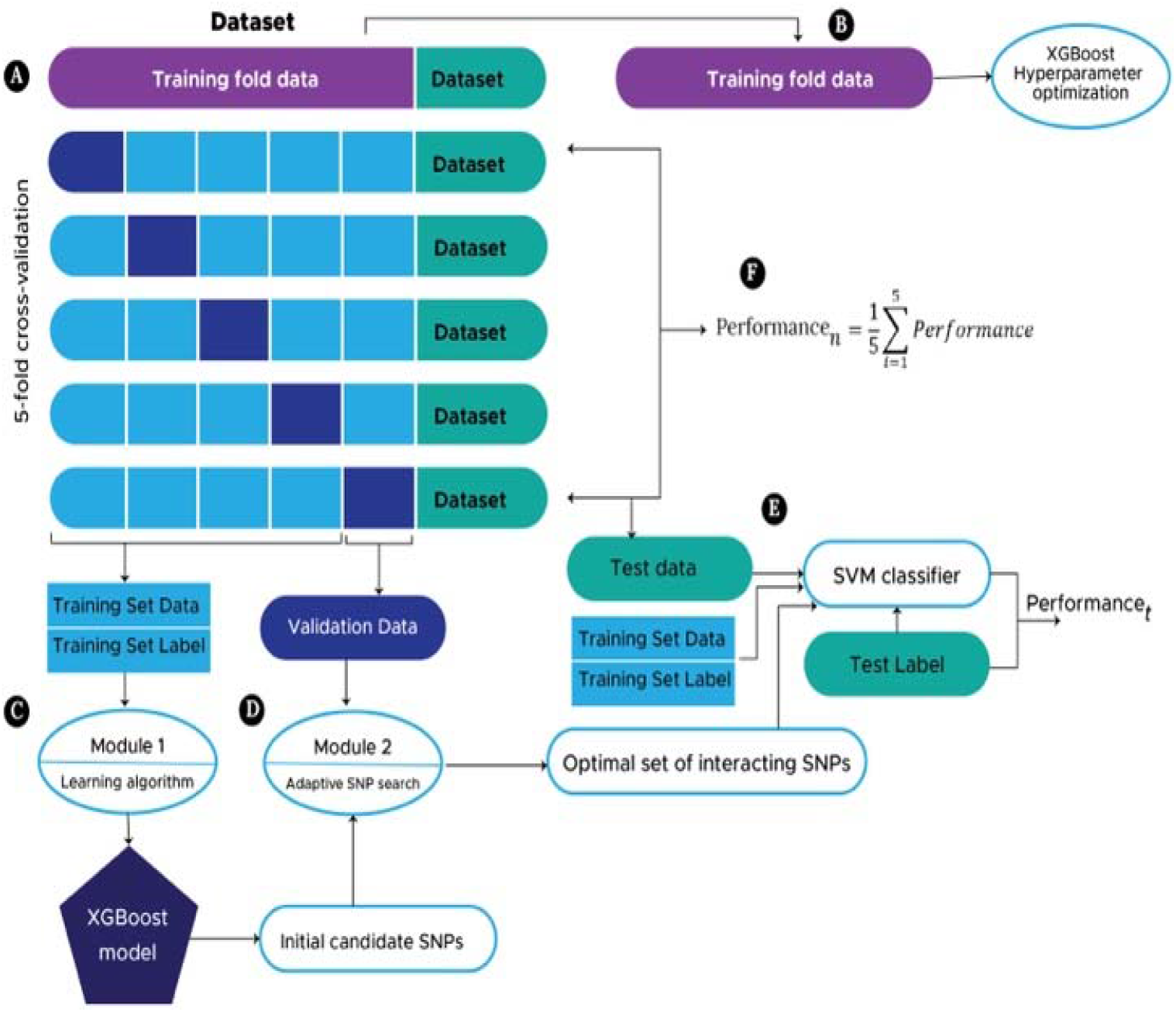
Algorithmic Framework SNP Selection Approach [5].

## 2. Method

### a. Data sourcing

We applied and executed the National Institute of Health (NIH) Database of Genotypes (DbGAP) protocols and requirements for obtaining licenses, file description, security and data sourcing. The study’s title is the Ghana Prostate Study [7](with dbGaP study accession number phs000838.v1.p1.).

The number of subjects consented to this study was 932 (474 prostate cancer cases and 458 controls). There are also data use limitations related to methods development research that align with the current methodological objectives. Whole genome genotyping was done on Illumina HumanOmni5-Quad with the identification of 4,301,332 SNPs.

### b. Quality Control

The Quality control was done in the protocol outlined by [1]. Plink (v 1.9) [8] R (4.03 version) was used for the research. The steps were as follows:

i. Delete SNPs and individuals with high levels of missingness
ii. Check for sex discrepancies and delete individuals with sex discrepancy
iii. Remove SNPs in the dataset that have a low minor allele frequency (MAF)
iv. Delete SNPs which are not in Hardy-Weinberg equilibrium (HWE)
v. And remove individuals with a heterozygosity rate deviating more than three standard deviations from the mean
vi. Check for population stratification using data from the 1000 Genomes Project.

### C. Perform Linear Association analysis

We Perform a genome-wide association testing approach for associating significant SNPs with the phenotype in a case-control GWAS. This is to identify SNPs that are significant with the regular GWAS. This would be compared with the top SNPs from the ML algorithm. In this logistic analysis, the logistic association used ten principal components from multidimensional scaling as covariates. Furthermore, we also do the chi-square association test with a Bonferroni corrected p-value, along with FDR and others. The previous two were done with the Plink software tool [8]. We also applied the linear genome-wide efficient mixed-model association using the Gemma tool [9].

### d. Optimize hyperparameters of XGBoost on the dataset

XGBoost is used to assess the significance of SNPs on a Prostate Cancer risk prediction job by offering a preliminary list of potential Prostate Cancer risk-predictive SNPs. We refer to this procedure as the first module of our suggested strategy. We utilized the gradient tree boosting method’s average of feature importance or gain. We leveraged the maximum power of XGBoost by tuning its hyperparameters. Hyperparameters optimized for the final feature importance scoring include Booster parameters: Learning rate, the full depth of a tree, subsample and learning task parameter: number of trees (n_estimators).

**Table 1:** Apply modified XGBoost algorithm on GWAS Data.

| S/N | Hyper-parameter optimizing space |  |  |
| --- | --- | --- | --- |
|  | n_estimators | max_depth | learning_rate |
| Set 1 | [50, 100] | [2, 4, 6, 8] | [0.001, 0.01, 0.1] |
| Set 2 | [150, 200] | [2, 4, 6, 8] | [0.001, 0.01, 0.1] |
|  | n_estimators | max_depth | learning_rate |
| Set 3 | [250, 300] | [2, 4, 6, 8] | [0.001, 0.01, 0.1] |

The modified learning algorithm is developed to find the optimal set of interacting SNPs with the best predictive capacity for predicting the genetic risk of developing prostate cancer.

The details of the algorithm are stated below in Fig 1 above:

1. Using a 4:1 proportion, divide the genotyped data into training fold and test data. A 5-fold stratified CV is used to partition further the training fold data: one-fold (validation data) is used to assess the set of identified SNPs generated by module 2, and the other four folds are combined to form a training set data for the XGBoost model, which is used to find initial candidate Prostate Cancer risk-predictive SNPs (module 1).
2. Using training fold data to optimize the XGBoost hyperparameters.
3. Module 1: Creating an initial list of possible Prostate Cancer risk-predictive SNPs using training set data to construct an XGBoost model
4. Module 2: Using the initial list of candidate SNPs obtained from C and the validation data, an adaptive iterative SNP selection method. The top interacting SNPs producing the best Prostate Cancer risk prediction accuracy on the validation data are chosen after SNPs are re-ranked (see Algorithm 1).
5. Using an SVM classifier, the top discovered interacting SNPs from (D) are adopted to forecast the Prostate Cancer risk on the test data.
6. Performance values are averaged across all trials to get the final accuracy in the test set.
7. The adaptive iterative SNP selection algorithm (Module 2) is as follows:
  a. Sort candidate SNPs from an XGBoost model in descending order.
  b. Choose M top- and M bottom-ranked. Separately re-rank M-selected SNPs from the bottom and top lists.
  c. Substitute the highest-/lowest-ranked SNP from the bottom/top with the lowest-/highest-ranked SNP from the top/bottom.
  d. Adaptively increase window size M by W
  e. Repeat 2 to 5 until the top and bottom lists overlap.
  f. Select the S top-ranked SNPs for the risk prediction model.

### e. Implementation details

We implement the approach with plink (1.9), xgboost (1.6.1), sci-kit-learn (1.1.1), matplotlib (3.5.2), pandas (1.4.3) and NumPy (1.23.1) all with python (3.8.10).

The platform was 72 CPUS (intel(r) Xeon (r) platinum 8124m CPU @ 3.00ghz) and a memory of 139 GB.

## 3. RESULTS

### a. Quality (QC) Control results’ visualization

After QC, the number of SNPs was 1,798,728. The results above show that the Quality control procedures were done effectively. These SNPs would be used as features for the ML algorithms. The number of samples was 907 after the completion. The plot below (Fig. 2) shows that after the QC and clustering, all our genotypes fell within the cluster of the African genomes. Plink-based MDS (multidimensional scaling) was used to provide ten principal components of genetic diversity for everyone, and this technique computes the genome-wide average fraction of alleles shared by every pair of people within the sample. They are then plotted to show where our samples fall using the 1000 genomes reference.

**Figure 2.**
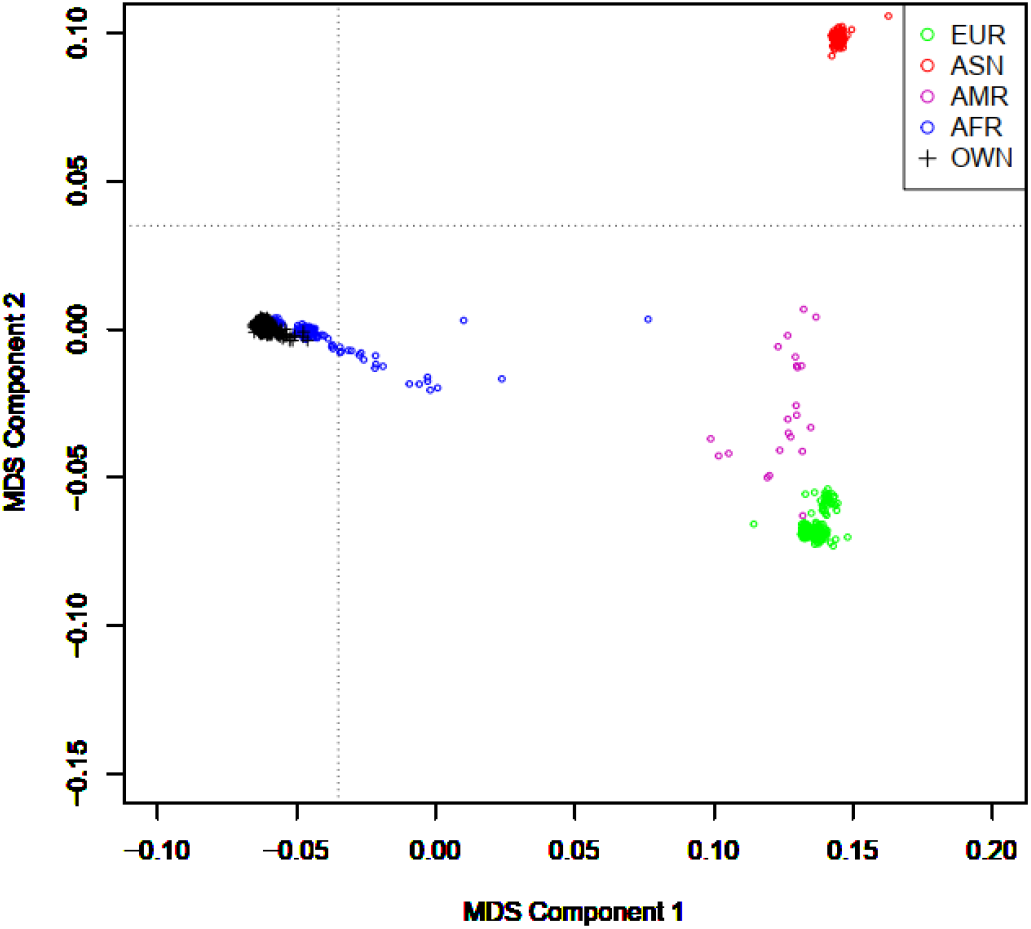
Population stratification of samples with the 1000 genomes (showing they are all African)

### b. Genome-wide statistical analysis

#### i. Logistic regression

Logistic regression is performed using the plink tool [8] and –logistic flag to test for the association of the SNPs with the binary outcome of having prostate cancer or not. The logistic regression model used the ten principal components calculated from the MDS step as covariates. Table II and Figure 3 below show the summary statics for the top 10 SNPs and quantile-quantile plots of the observed/expected log-transformed p-values.

**Table 2:** Logistic Regression Summary Statistics.

| *Chr* | *SNP* | *Allele 1* | *Odds Ratio* | *P-value* |
| --- | --- | --- | --- | --- |
| 6 | kgp6409239 | A | 0.4983 | 1.89E-06 |
| 13 | kgp12384188 | A | 0.5813 | 2.32E-06 |
| 6 | kgp1460575 | A | 0.5001 | 2.49E-06 |
| 3 | kgp3981243 | A | 0.6017 | 2.84E-06 |
| 12 | kgp2676687 | G | 1.603 | 3.75E-06 |
| 12 | rs7135604 | G | 1.603 | 3.75E-06 |
| 2 | rs875077 | G | 1.545 | 4.01E-06 |
| 1 | rs12057381 | A | 0.4608 | 5.03E-06 |
| 1 | kgp10577421 | A | 1.789 | 6.15E-06 |

**Figure 3.**
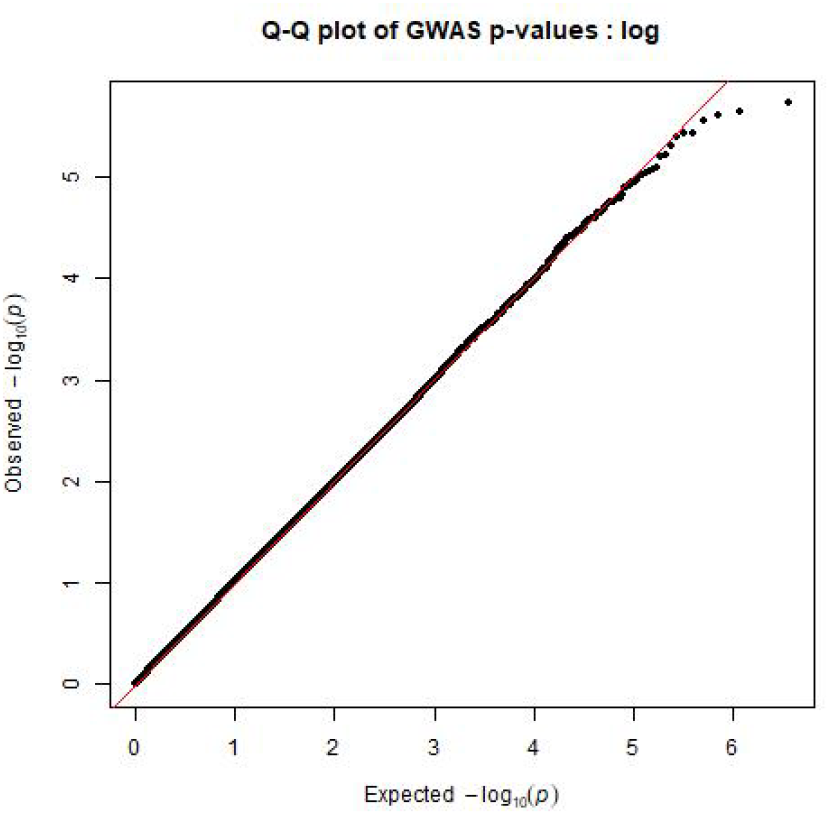
Q-Q plot of logistic association.

#### ii. Chi-square association analysis

The genomic inflation estimates (lambda) calculated for the genomic control analysis was 1.030 based on the median chi-square in the chi-square association analysis. Table II and Figure 4 show the summary statistics (including the chi-square statistic (CHISQ)) of the top ten SNPs and the quantile-quantile plots, respectively.

**Table 3:** Summary statistics of Chi-square association analysis.

| *Chr* | *SNP* | *P-value* | *Odds Ratio* | *CHISQ* |
| --- | --- | --- | --- | --- |
| 6 | kgp6409239 | 2.17E-07 | 0.4854 | 26.88 |
| 6 | kgp1460575 | 2.56E-07 | 0.4851 | 26.56 |
| 2 | rs875077 | 6.05E-07 | 1.606 | 24.9 |
| 12 | rs12302336 | 1.13E-06 | 2.134 | 23.69 |
| 1 | rs12057381 | 1.19E-06 | 0.4558 | 23.59 |
| 4 | rs6817894 | 1.87E-06 | 0.3918 | 22.72 |
| 11 | kgp9822254 | 2.11E-06 | 0.6265 | 22.5 |
| 18 | rs163050 | 2.24E-06 | 0.5952 | 22.38 |
| 10 | kgp2771589 | 2.86E-06 | 0.486 | 21.91 |

**Figure 4.**
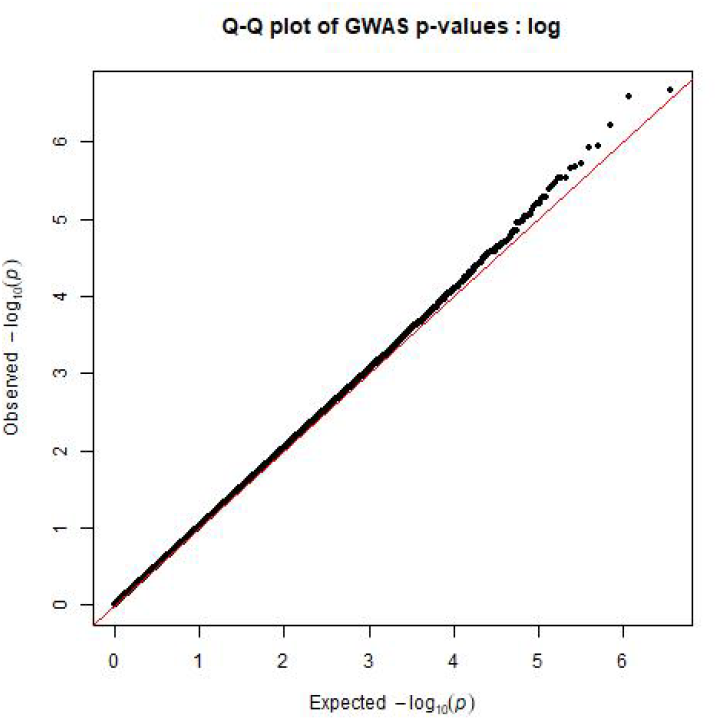
Q-Q plot of chi-square association analysis.

#### iii. Gemma Linear mixed model

We used the Gemma tool [9] to perform a univariate linear mixed model (LMM) implementation of the association testing. Table IV and Figure 5 below show the summary statics for the top 10 SNPs and quantile-quantile plots of the observed/expected log-transformed p-values, respectively.

**Table 4:** Summary Statistics of Linear Mixed Model.

| *Chr* | *SNP* | *Allele 1* | *Odds Ratio* | *P-value* |
| --- | --- | --- | --- | --- |
| 6 | kgp6409239 | A | -0.1652 | 4.45E-07 |
| 6 | kgp1460575 | A | -0.16444 | 5.74E-07 |
| 12 | rs12302336 | G | 0.17847 | 1.31E-06 |
| 1 | rs12057381 | A | -0.18354 | 1.63E-06 |
| 2 | rs875077 | G | 0.107467 | 1.68E-06 |
| 11 | kgp9822254 | A | -0.11431 | 2.31E-06 |
| 19 | rs9967612 | C | 0.116094 | 2.47E-06 |
| 18 | rs163050 | A | -0.12617 | 2.52E-06 |
| 2 | kgp12239910 | G | -0.11342 | 2.67E-06 |

**Figure 5.**
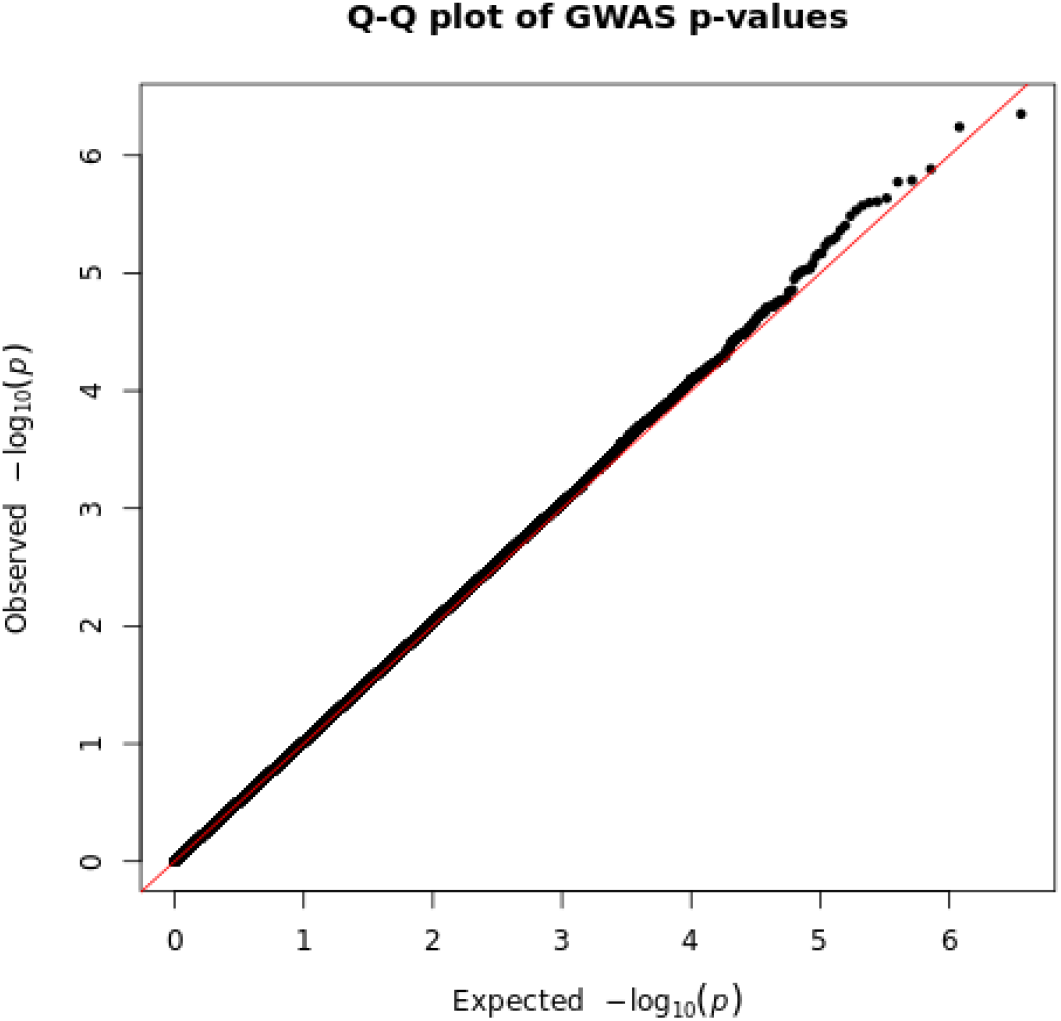
Q-Q plot of Linear mixed Models.

### c. XGBoost Model Evaluation

We tested our suggested approach using ten iterations of 5-fold cross-validation to address the issue of the inadequate number of genotyped BC data needed to build a highly performing Prostate cancer SVM risk prediction model (CV). There was an independent test set that we left out and to which the algorithm needed to be applied. The models were evaluated in the Training, Dev and Test set using AUC (Area under the curve), ROC (Receiver Operator Characteristic), Precision-Recall Curves and Mean Average Precision over the 50 folds. The ROC curve is a plot of the actual positive rate versus the false positive rate, and it evaluates machine learning models, and others listed heretofore.

#### i. Training Dataset

The risk-predictive SNPs discovered from the iterative SNPs selection approach were used to classify the trainset data. We performed ten repetitions of 5-fold cross-validation to get a perfect estimate of the model performance. The train’s Mean Average precision was 66.39 (with a standard error of 0.16). The AUC-ROC was 0.66, while the AUC under the Precision-Recall Curve was 0.69.

**Figure 6.**
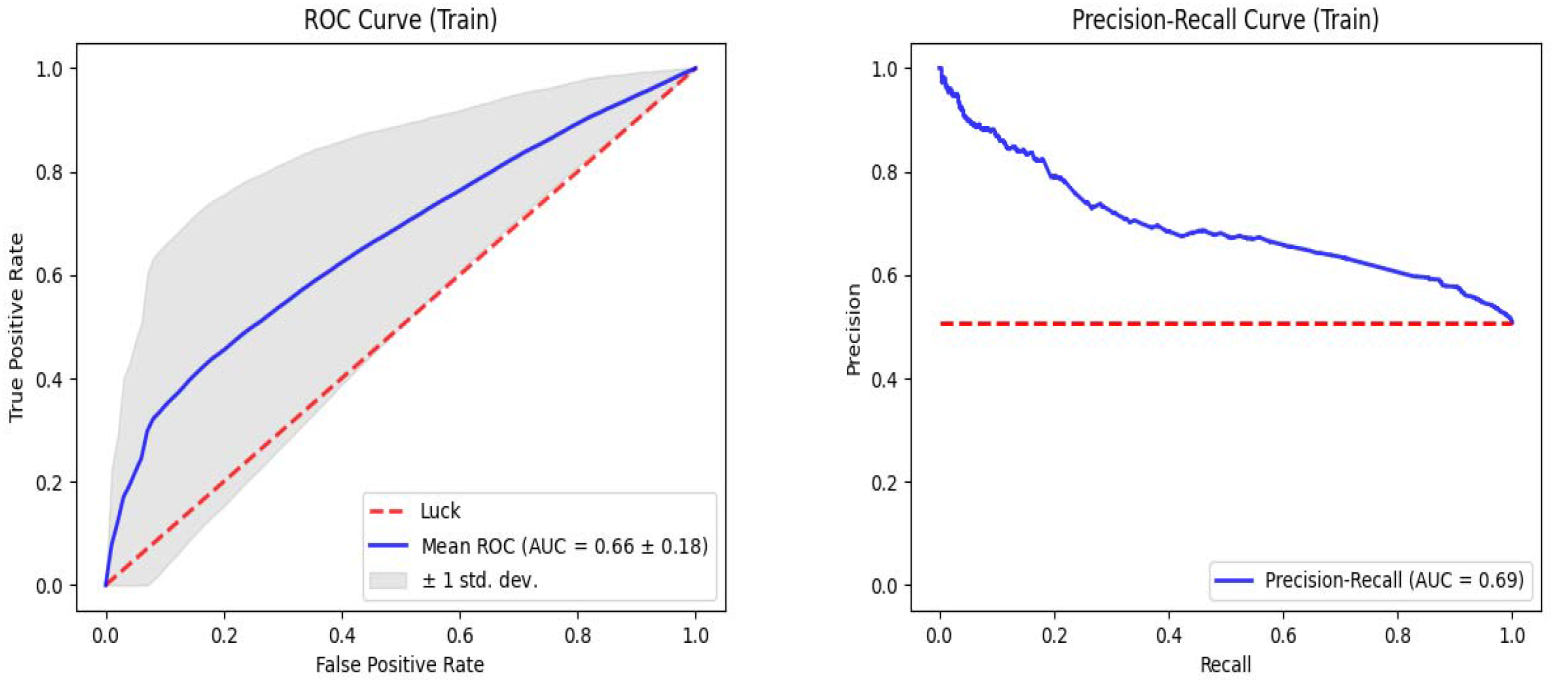
ROC curve of Training Set & The precision-recall curve of the Training set.

#### ii. Dev/Validation Dataset

In the Cross-validation approach, a Portion of the fivefold is used for validation, and this is used to ensure that the model does not overfit the training data. The validation Mean Average precision was 66.38 (with a standard error of 0.17). The AUC-ROC was 0.57, while the AUC under the Precision-Recall Curve was 0.60.

**Figure 7.**
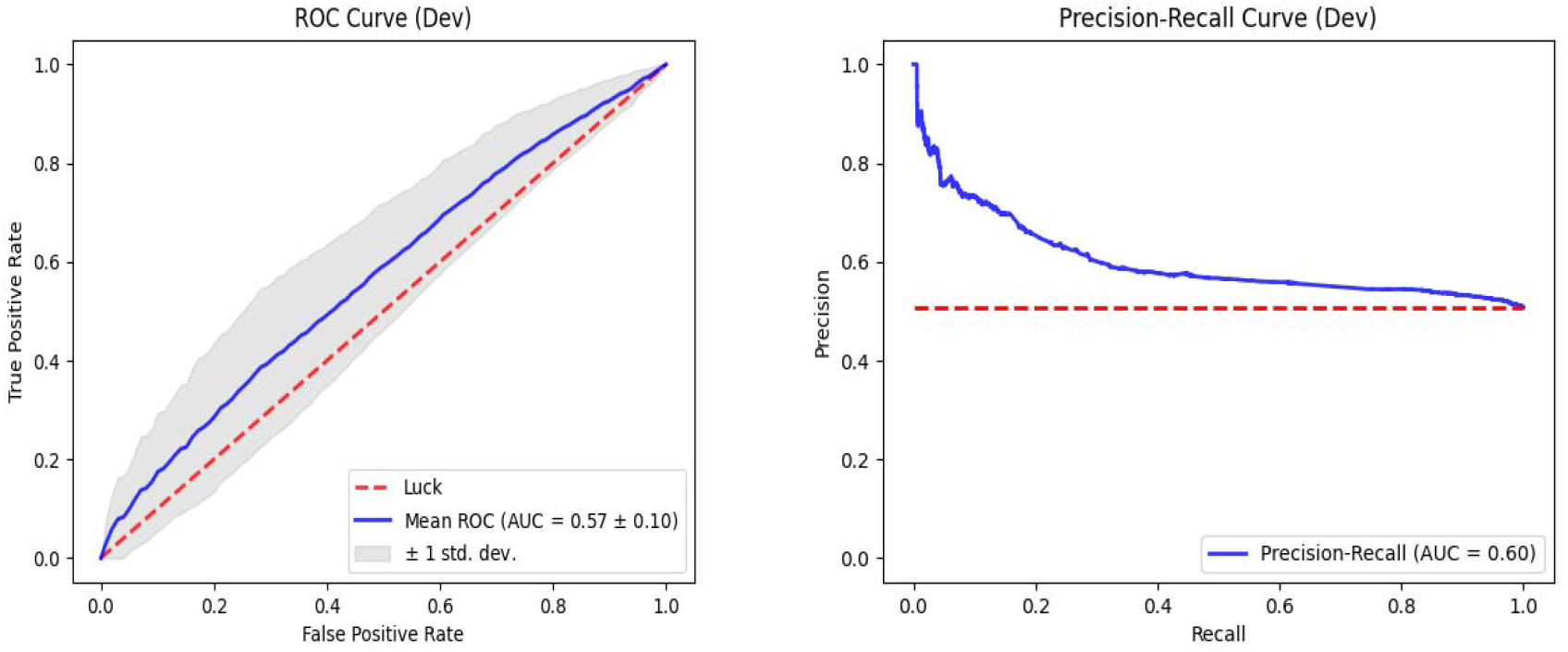
ROC curve of the Dev set & the precision-recall curve of the Dev set.

#### iii. Test Dataset

The test data is not used in the SNP selection process; instead, it is solely used to assess the final prediction accuracy of the SVM model. The test set Mean Average precision was 56.43 (with standard error: 0.07). The AUC-ROC was 0.55, while the AUC under the Precision-Recall Curve was 0.57.

**Figure 8.**
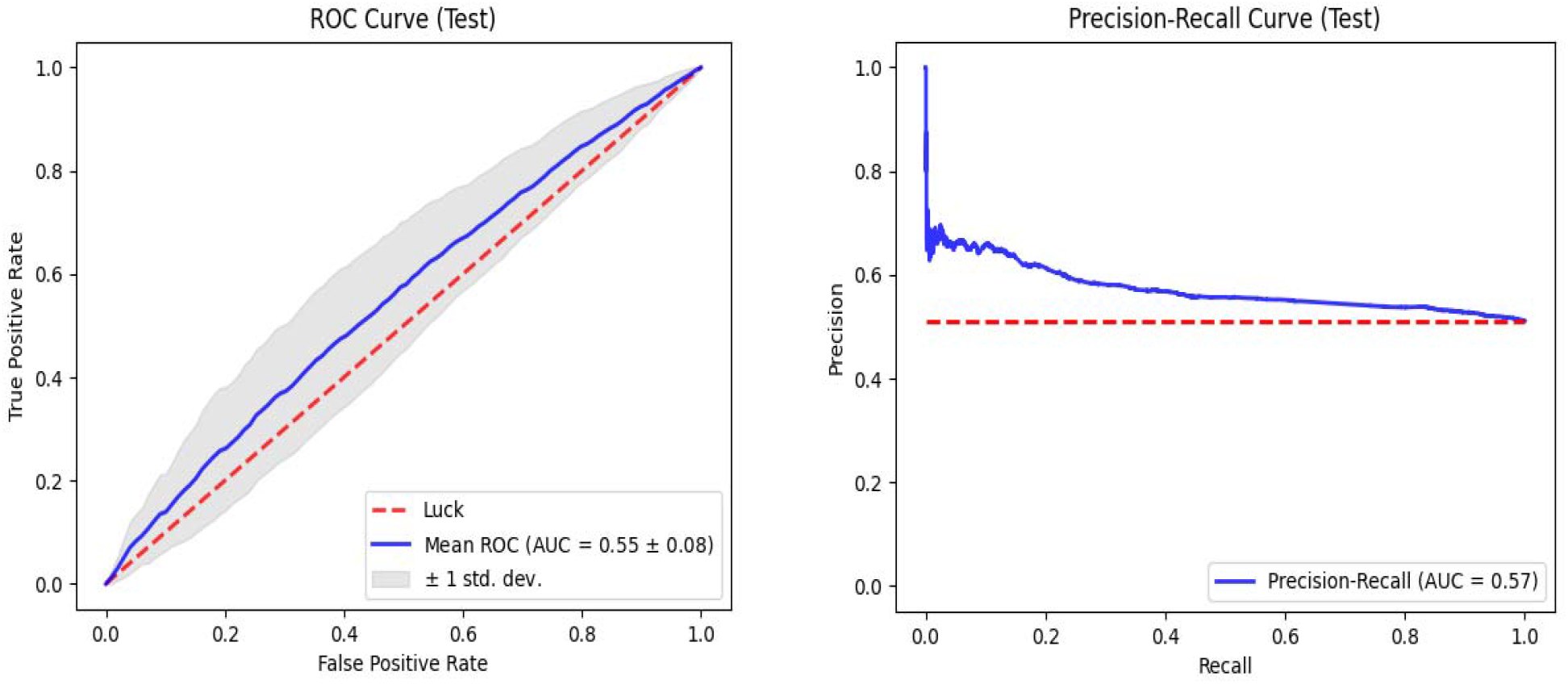
ROC curve of the Test set & the precision-recall curve of the Test set.

### d. Annotation of Top Risk Variants

After running the algorithm, the optimal set of SNPs on the classifier, which achieved the highest performance, was 2798 SNPs. These SNPs were then annotated with Ensembl (https://ensembl.org) to give a set of 847 genes associated with these SNPs. The 847 genes list were put into esyN (www.esyn.org) [10] to find these genes’ genetic and physical interactions. The database for the interaction was Homo Sapiens BioGrid. The number of Nodes was 339, and the number of edges was 622 in the interaction network. This showed that about 40% of these genes interact in the network. However, further delineating the interactions between the genes may be needed. This gives evidence of the interactions between the genes that were developed from the Machine Learning algorithm. It provides evidence that the XGBoost non-linear algorithms could identify interacting SNPs that affect the genetic trait in question, the development of prostate cancer.

**Figure 9.**
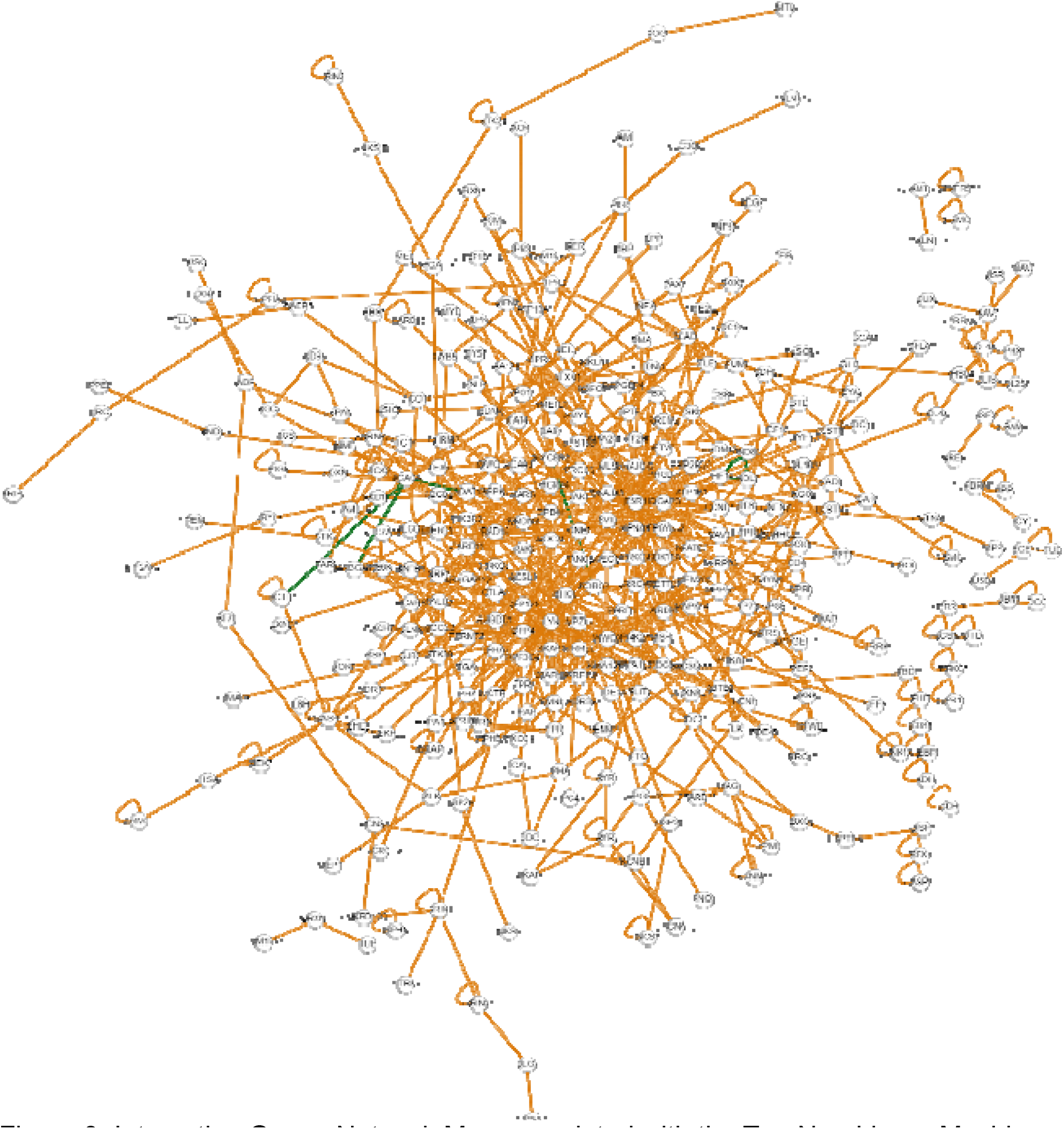
Interacting Genes Network Map associated with the Top Non-Linear Machine Learning Interacting SNPs.

After obtaining the summary statistics from the linear models, the summary statistics were filtered with the alpha-value (0.05/n). This is to give stringent p-value thresholding at p <5.5e-5. After which, an intersection operation was done between the machine learning model’s optimal interacting 2798 SNPs and the three linear models (Linear mixed, Adjusted association and logistic regression) to get the final 18 SNPs below.

SNPs and genes with “-”have no previously known reports in Ensembl, DbDSNP or ClinVar variant database. This shows that the results of the algorithm have some biological relevance. However, the number of intersection SNPs is expected to be low. This is because of the difference between the linear and non-linear approaches. Furthermore, functional enrichment was done with the set of variants to identify the citations bearing those genes in the PubMed database [11] using the G2SD tool. The results are seen in Table VI below.

**Table 5:**
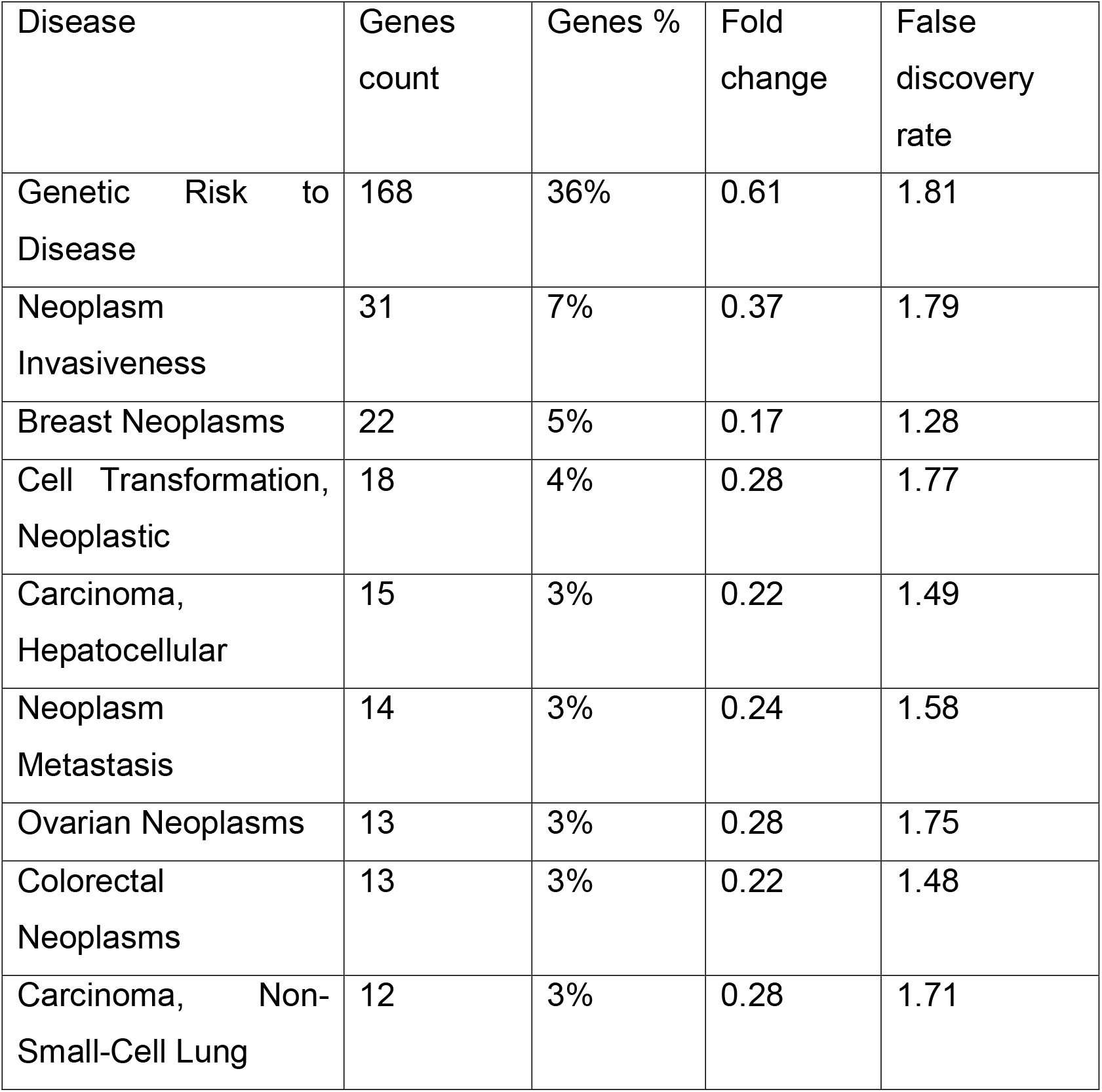
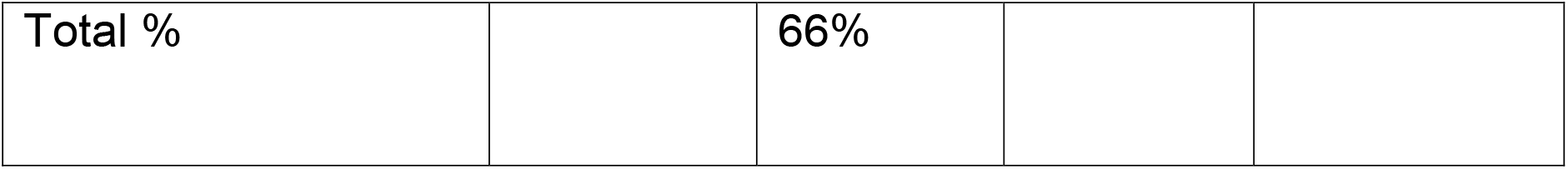
Rank Genes by Disease using Annotations of their PubMed Citations.

**Figure 10.**
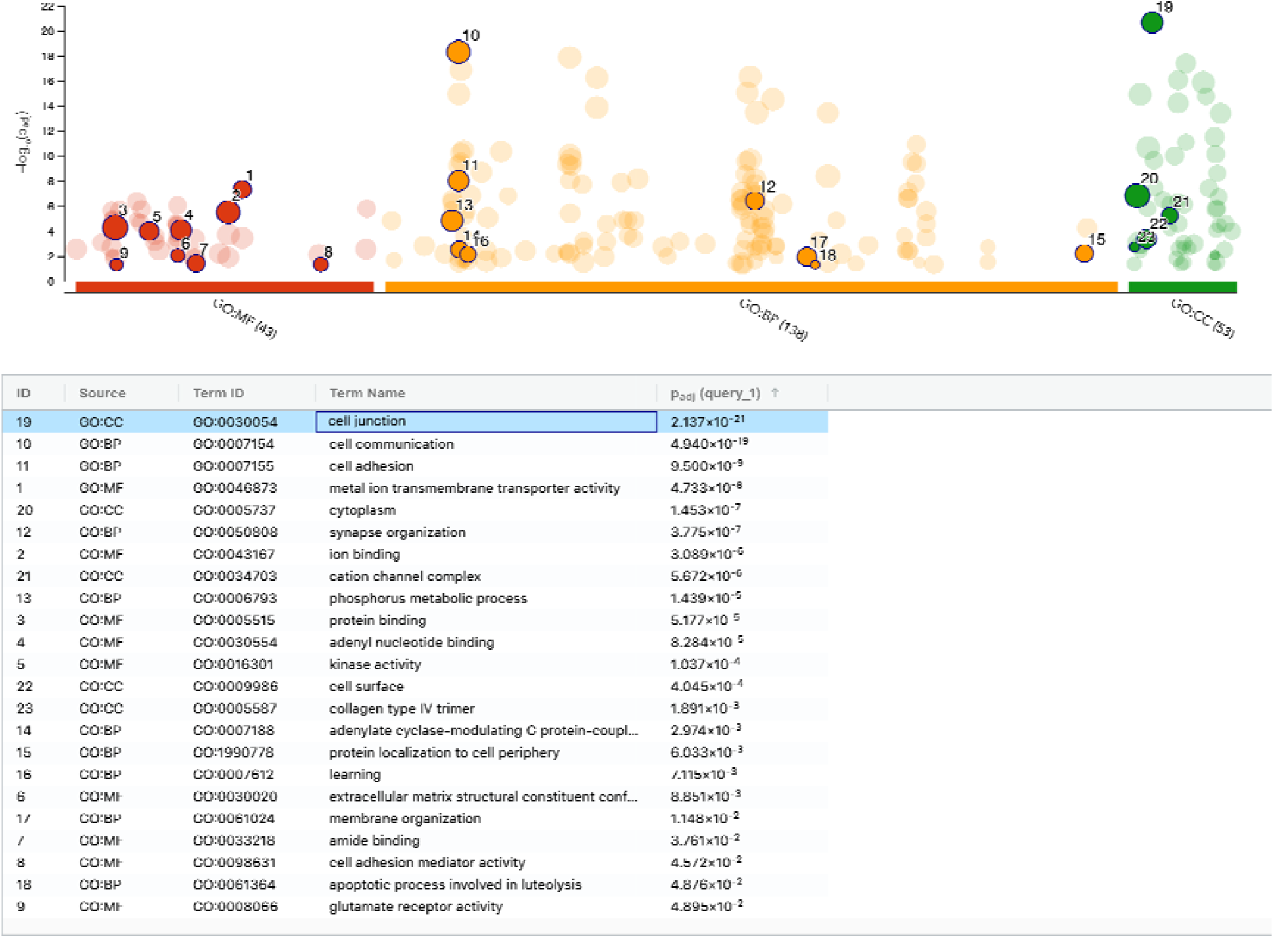
Gene ontology (GO) enrichment of the top Associated Genes.

**Figure 11.**
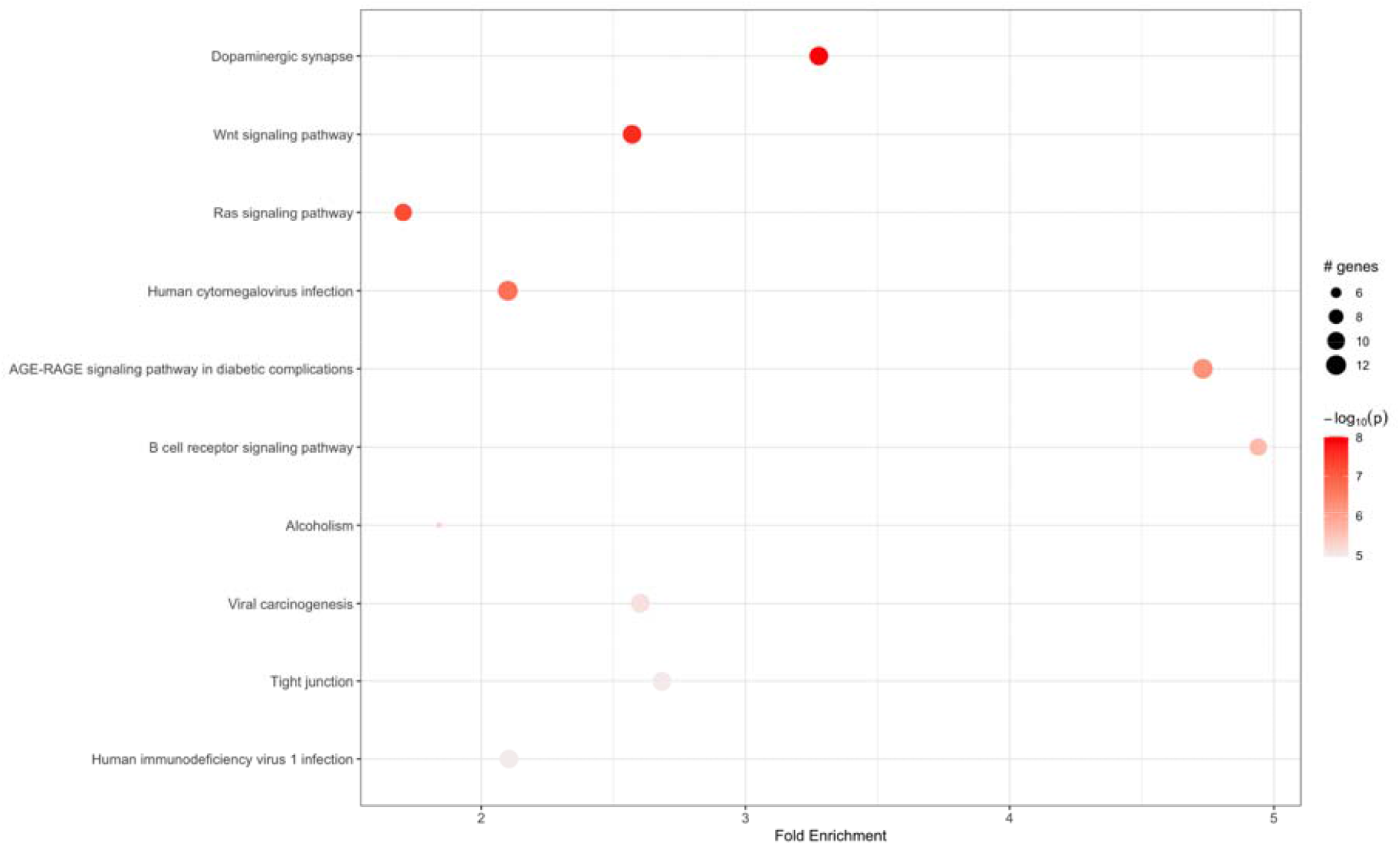
KEGG Pathway analysis of the top associated pathways.

**Figure 12.**
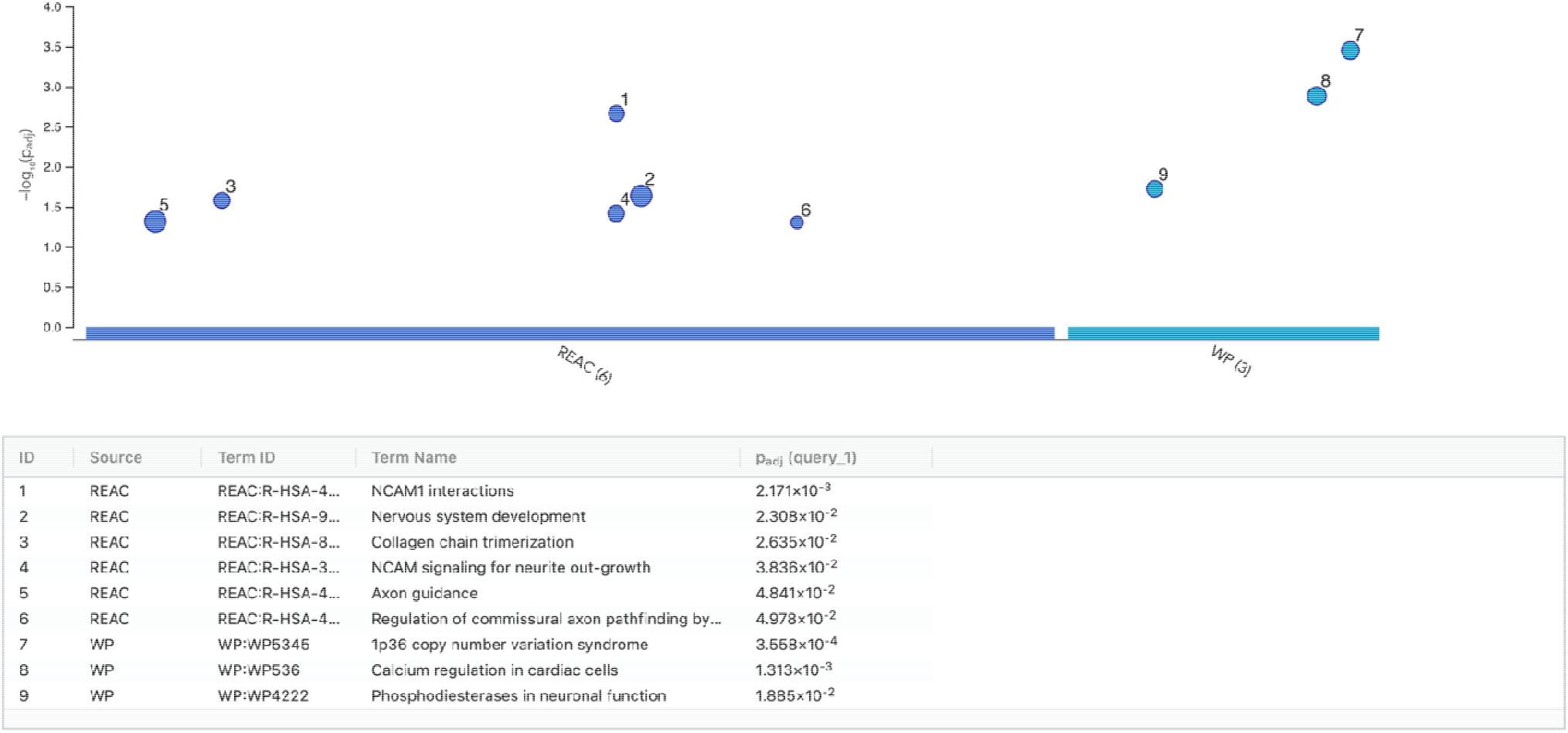
Reactome pathway enrichment of the top associated genes.

## 4. DISCUSSION

Limitations of traditional genome-wide association studies lie in the fact that they are not well suited to capture the non-linear effects of genetic variants on traits, such as the risk for developing prostate cancer, in this study, for instance. We hope to show in this work that non-linear approaches are better suited to complex traits such as prostate cancer disease biology. In contrast to traditional regression approaches, XGBoost adds complicated non-linear feature interactions into prediction models in a non-additive way. It has achieved cutting-edge results in several Kaggle (https://www.kaggle.com/) machine learning tasks. In previous work, [3] reviewed the approaches of ensembling algorithms in GWAS data. In this study, we had a sample size of 907, and we split it into 80% training, 10% test and 10% validation sets. However, the number of features was 1,798,727; we applied the algorithm in Figure 1. The adaptive iterative SNP search algorithms were done to get the optimal set of SNPs with the best performance in the SVM classification task.

Consequently, the train set had a Mean average precision of 66% after ten repetitions of 5-fold cross-validation on the test set. Furthermore, we achieved 69% for the area under the precision-recall curve. However, both values dropped to 56% and 57% on the test set. This was expected as the sample sizes dropped significantly after splitting into the test, validation and training sets. It would be helpful to increase the sample sizes in future work to improve the model for future work.

## 5. CONCLUSION

It is helpful to remember the sequence of computations: To predict Prostate Cancer risk using the identified interacting SNPs and an SVM classifier, the XGBoost model must first be trained. This involves obtaining initial candidates of Prostate Cancer risk-predictive SNPs, performing an adaptive iterative search over the candidate SNPs, and capturing the group of interacting SNPs with the highest Prostate Cancer risk-predictive potential.

Given the complex cancers, the non-additive interactions between genetic variants can explain the missing heritability [2]. Moreover, it has been shown that extreme gradient boosting algorithms can uncover interaction effects between several SNPs. These interactions occur, given the results above. This work aimed to apply a non-linear extreme gradient boosting machine learning algorithm to identify a set of SNPs (genetic components) that interact to affect the development of Prostate Cancer disease in an African population.

## Supporting information

Unique Gene list

Covariates for logistic regression analysis

Coordinates from KGP to RSIS

Unique Gene list-OPTIMAL SUBSET

## 6. Acknowledgement

The support of the Covenant Applied Informatics and Communication-African Centre of Excellence is Acknowledged for funding this work.

## Notes

https://github.com/davidenoma/prostate_cancer_genetic_association_risk_pred

### Competing Interest Statement

The authors have declared no competing interest.

